# Convergent mosaic brain evolution is associated with the evolution of novel electrosensory systems in teleost fishes

**DOI:** 10.1101/2021.09.29.462233

**Authors:** Erika L. Schumacher, Bruce A. Carlson

## Abstract

Brain region size generally scales allometrically with total brain size, but mosaic shifts in brain region size independent of brain size have been found in several lineages and may be related to the evolution of behavioral novelty. African weakly electric fishes (Mormyroidea) evolved a mosaically enlarged cerebellum and hindbrain, yet the relationship to their behaviorally novel electrosensory system remains unclear. We addressed this by studying South American weakly electric fishes (Gymnotiformes) and weakly electric catfishes (*Synodontis* spp.), which evolved varying aspects of electrosensory systems, independent of mormyroids. If the mormyroid mosaic increases are related to evolving an electrosensory system, we should find similar mosaic shifts in gymnotiforms and *Synodontis*. Using micro-computed tomography scans, we quantified brain region scaling for multiple electrogenic, electroreceptive, and non-electrosensing species. We found mosaic increases in cerebellum in all three electrogenic lineages relative to non-electric lineages and mosaic increases in torus semicircularis and hindbrain associated with the evolution of electrogenesis and electroreceptor type. These results show that evolving novel electrosensory systems is repeatedly and independently associated with changes in the sizes of individual brain regions independent of brain size, which suggests that selection can impact structural brain composition to favor specific regions involved in novel behaviors.

## Introduction

Brains are composed of multiple regions that vary widely in size across vertebrates and are associated with particular functions and behaviors (Striedter, 2005; Striedter and Northcutt, 2020). Much of the variation in brain region sizes is attributed to the allometric scaling of each region with total brain size (concerted evolution), which may result from conservation and constraint in developmental neurogenesis (Finlay and Darlington, 1995; Striedter, 2005). Seemingly disproportionately enlarged regions can instead have larger allometric slopes by extending the timing of neurogenesis for late developing brain regions such as the cortex and cerebellum (Finlay and Darlington, 1995). However, changes in brain region sizes independent of total brain size, or mosaic shifts, have also been observed in several taxa and are hypothesized to reflect selection on traits associated with those regions (Barton and Harvey, 2000; Striedter, 2005). Mosaic shifts in fine-scale brain regions and circuits have been linked to changes in behavior (Carlson et al., 2011; Vélez et al., 2019; Gutiérrez-Ibáñez et al., 2014; Moore and DeVoogd, 2017; DeCasien and Higham, 2019; Krebs, 1990), but the scale at which selection can act to alter brain region sizes remains unclear. Most studies looking at major brain regions instead find evidence of concerted evolution (Finlay and Darlington, 1995; Striedter, 2005; Yopak et al., 2010). There is some evidence of mosaic evolution at these scales (Hoops et al., 2017; Sukhum et al., 2018), but the drivers and selective pressures necessary for mosaic evolution to overcome constraints remain unclear. Further, this is difficult to test without repeated evolution of the same phenotypes.

Mosaic brain evolution of major brain regions is hypothesized to occur more frequently at larger taxonomic scales and alongside behavioral innovations that open new niches since mosaic shifts are more likely to contribute to major differences in brain function (Striedter, 2005). In dragon lizards, mosaic brain evolution is associated with ecomorph (Hoops et al., 2017), but as many different behavioral and sensory changes occur alongside ecomorph development, it is difficult to identify specific selective pressures favoring the observed ecomorph brain structure. Weakly electric fishes are excellent for testing whether mosaic brain evolution occurs with behavioral novelty: these fishes evolved behaviorally novel active electrosensory systems with several neural innovations, which likely resulted in strong selection for electrosensory processing capabilities (Carlson and Arnegard, 2011). Further, multiple lineages independently evolved similar electrosensory systems (Crampton, 2019). Previous studies found that African weakly electric fishes (Mormyroidea) evolved extremely large brains along with mosaic increases in the sizes of the cerebellum and hindbrain relative to other non-electric osteoglossiforms (Sukhum et al., 2018, 2016). These mosaic increases occurred alongside the evolution of an active electrosensory system (electrogenesis + electroreception), but since this is only a single lineage, it is impossible to determine whether these mosaic shifts are associated with the evolution of this electrosensory system or with other phenotypes that differentiate mormyroids from their closest living relatives. Further, one non-electric osteoglossiform, *Xenomystus nigri*, is electroreceptive, and there is no evidence that it has experienced mosaic brain evolution compared to other non-electric osteoglossiforms.

Here, we investigated another lineage of fishes, otophysans, which includes taxa that evolved a similar electrosensory system to varying degrees independent of mormyroids, to determine whether these mosaic shifts are found repeatedly alongside the evolution of active electrosensing. Although osteoglossiform and otophysan lineages have convergently evolved similar electrosensory systems (Crampton, 2019), there are some distinctive differences in the degree of electrosensory usage, electroreceptor type, and electrical discharge type that could indicate differential selective pressures on the brains of these species. These differences allowed us to assess how multiple aspects of electrosensory systems relate to mosaic brain evolution.

## Results

### Electrogenic species have similar structural brain variation

To investigate how electrosensory systems relate to structural brain variation, we combined published osteoglossiform data for electrogenic mormyroids, electroreceptive *Xenomystus*, and non-electrosensory outgroup species (Sukhum et al., 2018) with otophysan data for two additional electrogenic lineages (Gymnotiformes and *Synodontis* Siluriformes), electroreceptive (but not electrogenic) siluriforms, and non-electrosensory Characiformes and Cypriniformes (outgroup otophysans). Using micro-computed tomography (μCT) scans, we measured total brain and brain region volumes for 15 electrogenic, 3 electroreceptive, and 4 non-electrosensory otophysan species. Combined with the published osteoglossiform data, this yielded a dataset of 32 species (Figure 1, Figure 1—figure supplement 1). We measured the volumes of seven distinct brain regions (olfactory bulbs, OB; telencephalon, TEL; hindbrain, HB; optic tectum, OT; torus semicircularis, TS; cerebellum, CB; and rest of brain, RoB) to determine patterns of major brain region scaling across taxa. Rest of brain includes thalamus, hypothalamus, and additional midbrain structures excluding optic tectum and torus semicircularis (see Materials and Methods, Brain Region Delimitation). To determine if the evolution of an electrosensory system is associated with extreme encephalization, we directly compared relative brain size residuals using a single, phylogenetic generalized least squares (PGLS) regression fit across all taxa. Considering the phylogenetic relationships among otophysans are still debated (Crampton, 2019; Hughes et al., 2018; Rabosky et al., 2013), our goal was only to account for relatedness to the best of our ability, not to propose a resolved phylogeny. We found that brain size varies in electrosensory and non-electrosensory otophysans but that gymnotiform and siluriform brains are not as enlarged as mormyroid brains, which suggests that extreme increases in brain size are not directly tied to electrosensory system evolution (Figure 2).

**Figure 1.**
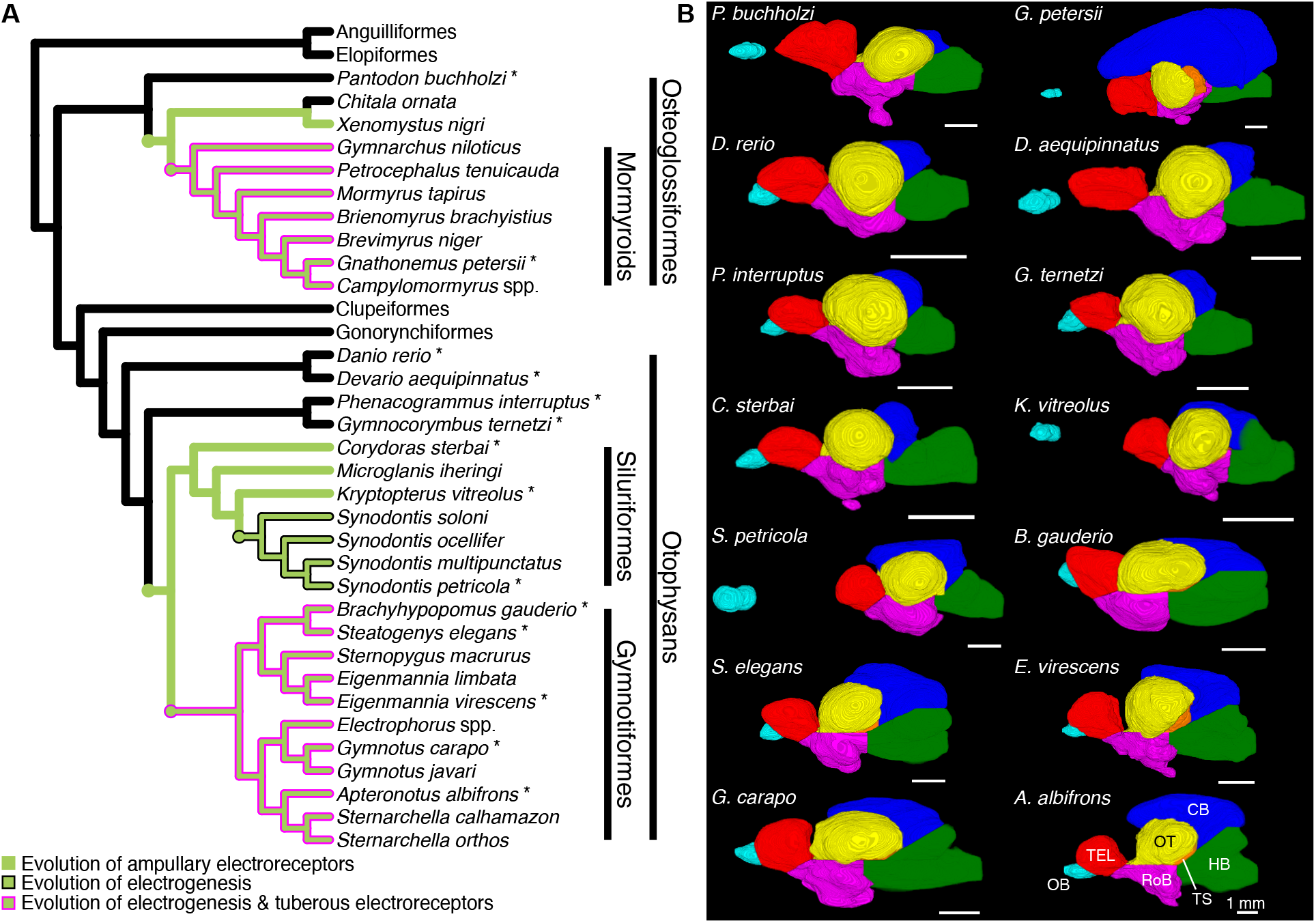
Brain morphology varies across species. (A) Cladogram of the inferred phylogenetic relationships of species included in this study (N=32) and the orders between them. Order level relationships are based on (Hughes et al., 2018). Green branches represent presence of ampullary electroreceptors. Black outline represents electrogenic species while the magenta outline represents electrogenic species with tuberous electroreceptors. (B) Example 3D reconstructions of brains from this study; these species are indicated on the cladogram with an asterisk. Brains are oriented from a lateral view with anterior to the left and dorsal at the top. Brain regions are color coded: OB = olfactory bulbs (cyan), TEL = telencephalon (red), HB = hindbrain (green), OT = optic tectum (yellow), TS = torus semicircularis (orange), CB = cerebellum (blue), RoB = rest of brain (magenta). Scale bar = 1 mm. **Figure supplement 1.** Video of rotating 3D reconstructions.

**Figure 2.**
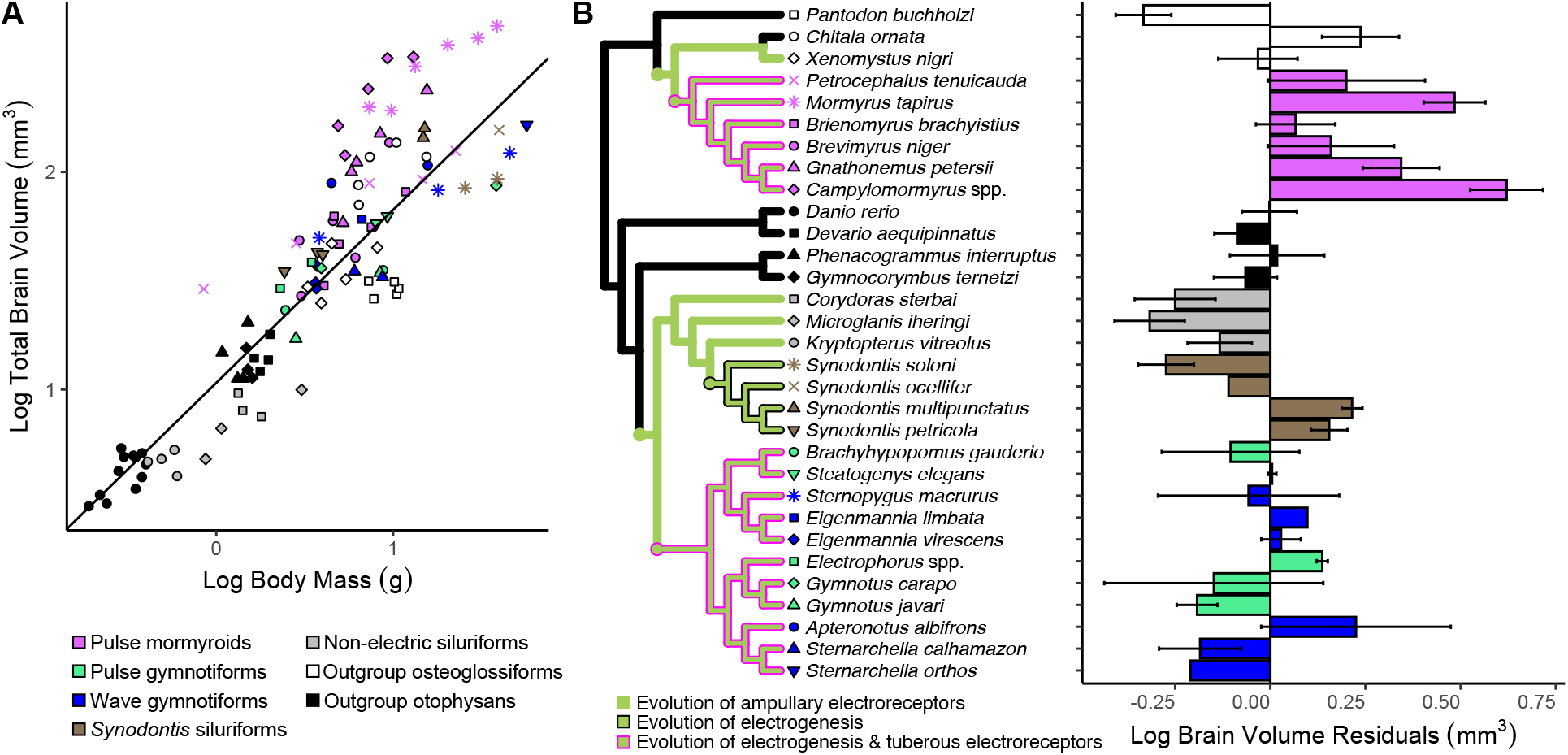
Mormyroids are more encephalized than gymnotiforms. (A) Log total brain volume by log body mass plot. Each point represents an individual, shapes correspond to species, and colors correspond to lineages: pink = pulse mormyroids (N=6), white = outgroup osteoglossiforms (N=3), blue = wave gymnotiforms (N=6), green = pulse gymnotiforms (N=5), brown = *Synodontis* siluriforms (N=4), grey = non-electric siluriforms (N=3), black = outgroup otophysans (N=4). PGLS line was determined from species means. (B) Average brain size residuals for each species calculated from A and organized phylogenetically. Error bars indicate standard deviation. Green branches on the cladogram represent presence of ampullary electroreceptors; black outline represents electrogenic species while the magenta outline represents electrogenic species with tuberous electroreceptors.

To determine how brain structure varies in association with electrosensory phenotype, we used regional measurements for all taxa and ran a phylogenetically corrected principle components analysis (pPCA). We found that electrogenic species cluster distinctly from both electroreceptive and non-electrosensory species, which overlap considerably (Figure 3A). All electroreceptive lineages (mormyroids, *Xenomystus*, gymnotiforms, and siluriforms) have evolved ampullary electroreceptors, which detect relatively low frequency electrical information, while only gymnotiforms and mormyroids have evolved additional tuberous electroreceptors that broaden the frequency range of detectable signals (Crampton, 2019). We find that electrogenic species with both electroreceptor types cluster distinctly from electrogenic fishes with only ampullary electroreceptors (*Synodontis* siluriforms). The first principle component (PC1) explained 92.02% of the variation in brain region volumes and is strongly correlated with total brain volume (ρ = −0.99, p < 10^-16^). PC2 explained 4.81% of the total variation, which is 60.3% of the variation in region volumes not explained by total brain volume. Whereas all of the brain regions load in the same direction for PC1, electrosensory associated cerebellum, torus semicircularis, and hindbrain load negatively on PC2 while the remaining regions (telencephalon, rest of brain, optic tectum, and olfactory bulbs) load positively. This suggests that concerted brain evolution explains the most variation in region volumes as seen in PC1, but that mosaic brain evolution could be contributing to the observed variation in brain region volumes as seen in PC2.

**Figure 3.**
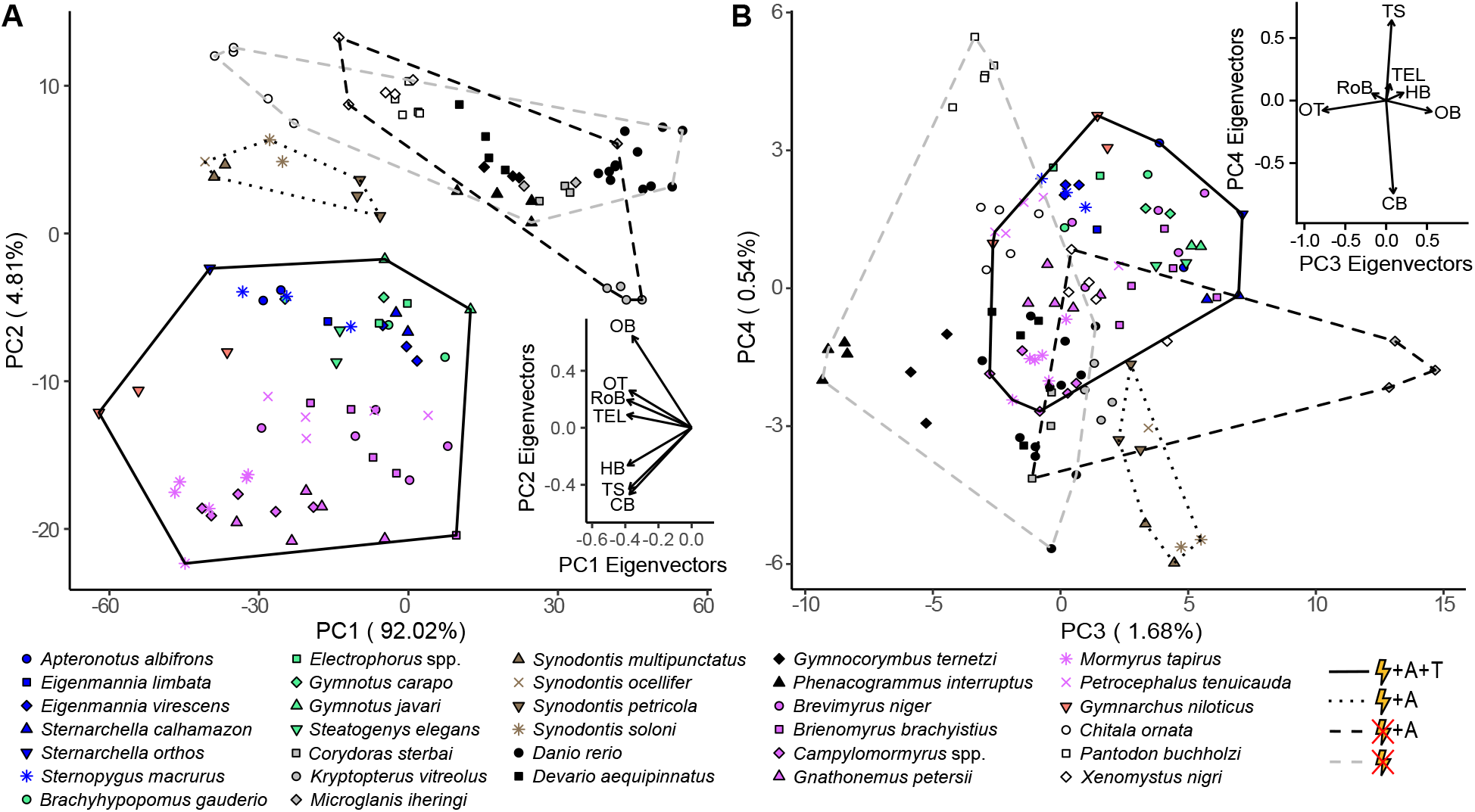
Species cluster distinctly in PC space based on electrosensory phenotype with electrosensory associated brain regions (HB, TS, CB) loading in the direction of electrogenic taxa for PC1 and PC2 (A), but not for PC3 and PC4 (B). Each point represents an individual, shapes correspond to species, and colors correspond to lineages: orange = wave mormyroid (N = 1), pink = pulse mormyroids (N = 6), white = outgroup osteoglossiforms (N = 3), blue = wave gymnotiforms (N = 6), green = pulse gymnotiforms (N = 5), brown =*Synodontis* siluriforms (N = 4), grey = non-electric siluriforms (N = 3), black = outgroup otophysans (N = 4). Minimum convex hulls correspond to electrosensory phenotypes: electrogenic + ampullary + tuberous electroreceptors (solid), electrogenic + only ampullary electroreceptors (dotted), only ampullary electroreceptors (black dashed), and non-electrosensory (grey dashed). Insets shows PC eigenvectors of each brain region. OB = olfactory bulbs, TEL = telencephalon, HB = hindbrain, OT = optic tectum, TS = torus semicircularis, CB = cerebellum, RoB = rest of brain. **Figure supplement 1.** pPCA model selection results.

To assess the relative importance of electrosensory phenotypes in explaining the axes of brain structural variation (PCs 1-4), we ran candidate models that considered body mass, total brain volume, presence or absence of electrogenesis, and electroreceptor type (tuberous and ampullary vs only ampullary vs none). Models that only consider allometric scaling with body mass and total brain volume would be consistent with concerted evolution while models that also consider either one or both electrosensory phenotypes would be in line with mosaic brain evolution since more than just allometric scaling explains the observed variation in brain region volumes. Since PC1 strongly correlates with brain size, we removed total brain volume as a variable from all PC1 models. We found that the model that considers the electrogenesis phenotype better explained PC1, but is statistically indistinguishable from the concerted model (Figure 3—figure supplement 1), which further supports the role of concerted evolution in determining the sizes of individual brain regions. The model that considers electroreceptor type better explained PC2 (Figure 3—figure supplement 1), which supports our hypothesis that the electrosensory system is related to mosaic evolutionary changes in brain region scaling. PC3 explains 1.68% of the total variation (21.1% of the variation not explained by total brain volume) and largely reflects the variation between olfactory bulbs and optic tectum with no separation between electrosensory phenotypes (Figure 3B). The model that considers electroreceptor type better predicts PC3 but is statistically indistinguishable from the concerted model, suggesting that this axis of brain variation likely evolved concertedly with brain size (Figure 3—figure supplement 1). PC4 explains 0.54% of the total variation (6.8% of the variation not explained by total brain volume), largely reflects the variation between torus semicircularis and cerebellum, and is better predicted by the model that considers both electrosensory phenotypes (Figure 3B, Figure 3—figure supplement 1). Taken together, these results highlight the importance of both concerted and mosaic brain evolution in producing the observed variation in brain region volumes.

### Mosaic shifts in electrogenic species relative to non-electric species

To directly test for mosaic shifts associated with electrosensory phenotypes, we fit PGLS regressions for each brain region against total brain size for species with each electrosensory phenotype and used an analysis of covariation (ANCOVA) to test for significant differences in the PGLS relationships for each brain region. Considering the debate on how best to assess patterns of brain region scaling (Finlay and Darlington, 1995; Yopak et al., 2010), we also determined PGLS relationships for each brain region against total brain volume minus the focal brain region and for pairwise region by region comparisons (Figure 4—figure supplement 1,2). Our results were broadly consistent across each method, so we present here the results for PGLS relationships of each brain region against total brain volume.

Since electroreceptive only species and non-electrosensory species overlapped considerably in the pPCA, we combined all non-electric taxa and performed an ANCOVA with pairwise posthoc testing for electrogenic taxa with both tuberous and ampullary electroreceptors (mormyroids and gymnotiforms), electrogenic taxa with only ampullary electroreceptors (*Synodontis* siluriforms), and non-electric taxa (Figure 4, Figure 4—figure supplement 4). We found a significant increase in y-intercept in cerebellum for all electrogenic species relative to non-electric species (p < 0.05). For torus semicircularis, we found a significant increase in y-intercept in electrogenic + ampullary only taxa relative to non-electric taxa (p < 0.05) and a further increase in y-intercept in electrogenic + ampullary + tuberous taxa (p < 10^-4^). We found a significant increase in y-intercept in hindbrain for electrogenic + ampullary + tuberous taxa relative to non-electric taxa (p < 0.01) with electrogenic + ampullary only taxa being intermediate (p > 0.05).

**Figure 4.**
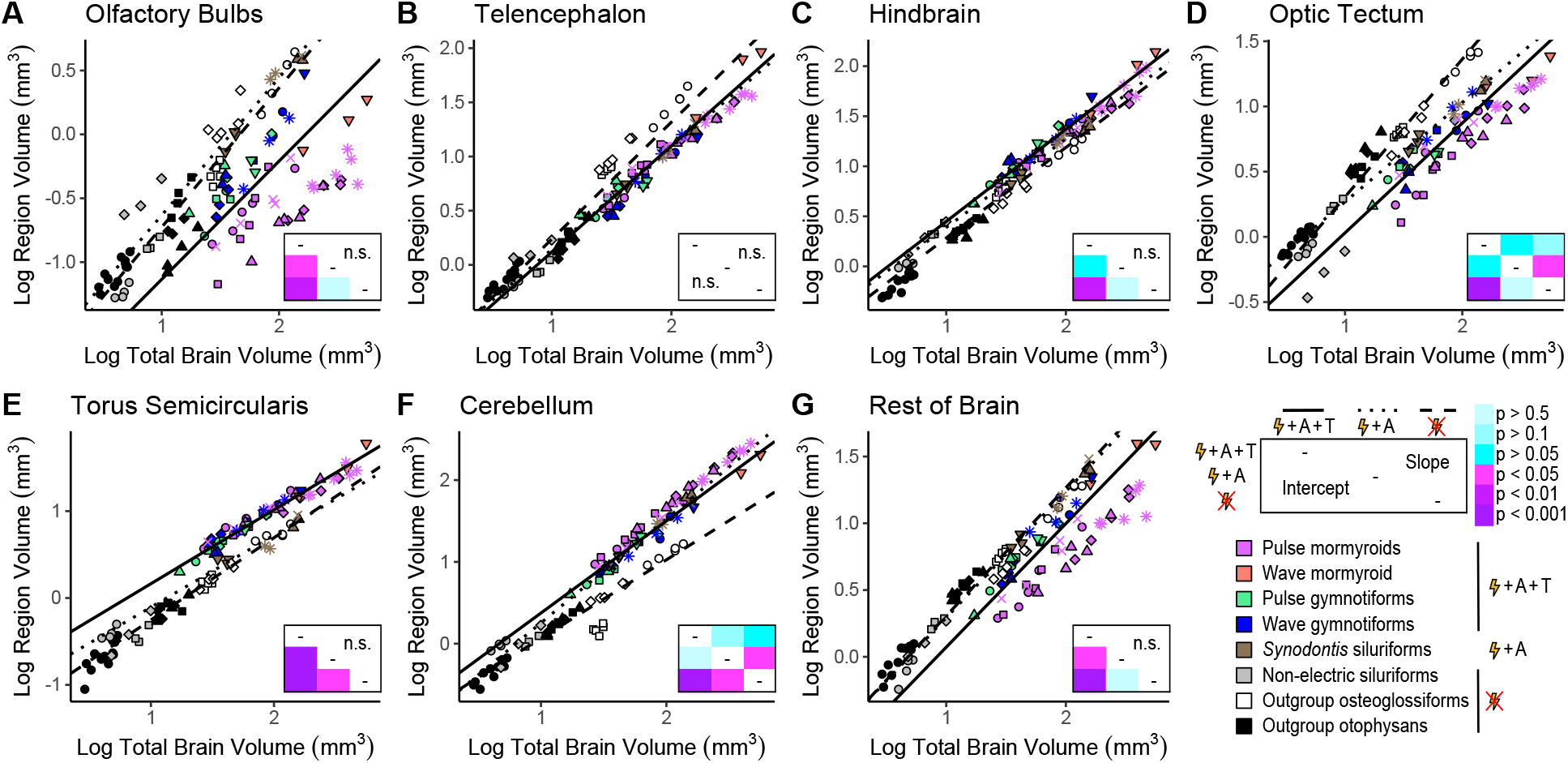
Mosaic increases in hindbrain, torus semicircularis, and cerebellum in electrogenic species with ampullary & tuberous electroreceptors. Plots of log region volume against log total brain volume for olfactory bulbs (A), telencephalon (B), hindbrain (C), optic tectum (D), torus semicircularis (E), cerebellum (F), and rest of brain (G). Each point corresponds to an individual and shapes represent the same species as Figure 3. PGLS lines were determined from species means and correspond to electrosensory phenotypes that cluster distinctly in Figure 3: electrogenic + ampullary + tuberous electroreceptors (pink, orange, green, blue points; solid line; N = 18), electrogenic + only ampullary electroreceptors (brown points; dotted line; N = 4), and non-electric (grey, white, black points; dashed line; N = 10). Inset shows a heatmap of the pairwise posthoc ANCOVA results with a Bonferroni correction for differences in intercept below the diagonal and differences in slope above the diagonal for each brain region. Significant differences are in shades of magenta/purple and non-significant differences are in shades of blue. **Figure supplement 1.** Similar mosaic shifts in comparisons of log region volume against log total brain – region volume. **Figure supplement 2.** Similar mosaic shifts in region by region comparisons. **Figure supplement 3.** No mosaic shifts found in electrogenic chondrichthyans. **Figure supplement 4.** ANCOVA results for electrosensory phenotype comparisons. **Figure supplement 5.** Pairwise posthoc ANCOVA results for electrosensory phenotype comparisons. **Figure supplement 6.** Effect size estimate of ANCOVA results for electrosensory phenotype comparisons.

There were significant decreases in y-intercept in olfactory bulbs and rest of brain for electrogenic + ampullary + tuberous taxa relative to both electrogenic + ampullary only and non-electric taxa (p_OB_ < 0.05, p_RoB_ < 0.05). For optic tectum, we found a significant decrease in y-intercept in electrogenic + ampullary + tuberous taxa relative to non-electric taxa (p < 10^-4^) with electrogenic + ampullary only taxa as intermediate (p > 0.05). There were no significant differences in y-intercept for telencephalon (p > 0.05). These results show that there are similar mosaic shifts in lineages that independently evolved electrogenesis. We also found a significant increase in slope in cerebellum and a significant decrease in slope in optic tectum for electrogenic + ampullary only species relative to non-electric species (p_CB_ < 0.05, p_OT_ < 0.05), which may be related to the reduced species sampling and brain size distribution of electrogenic + ampullary only species (N = 4) relative to non-electric (N = 10) and electrogenic + ampullary +tuberous species (N = 18).

### Lineage-specific mosaic shifts within electrosensory phenotypes

To determine if there are lineage-specific differences within electrosensory phenotypes, we performed an ANCOVA of each brain region between mormyroids and gymnotiforms (Figure 5A, Figure 5—figure supplement 1) and between non-electric osteoglossiforms and otophysans (Figure 5B, Figure 5—figure supplement 1). We found a shallower slope and smaller y-intercept for mormyroids in olfactory bulbs (p_slope_ < 10^-4^, p_intercept_ < 0.05) and rest of brain (p_slope_ < 10^-2^, p_intercept_ < 0.05), shallower slope for mormyroids in torus semicircularis (p < 10^-2^), and steeper slope for mormyroids in cerebellum (p < 0.05). These differences are likely contributing to the secondary clustering between mormyroids and gymnotiforms in the pPCA (Figure 3A). For non-electric fishes, we found that osteoglossiforms have a significantly larger y-intercept for telencephalon (p_TEL_ < 10^-2^), but no differences in either slope or y-intercept in the other brain regions (p > 0.1).

**Figure 5.**
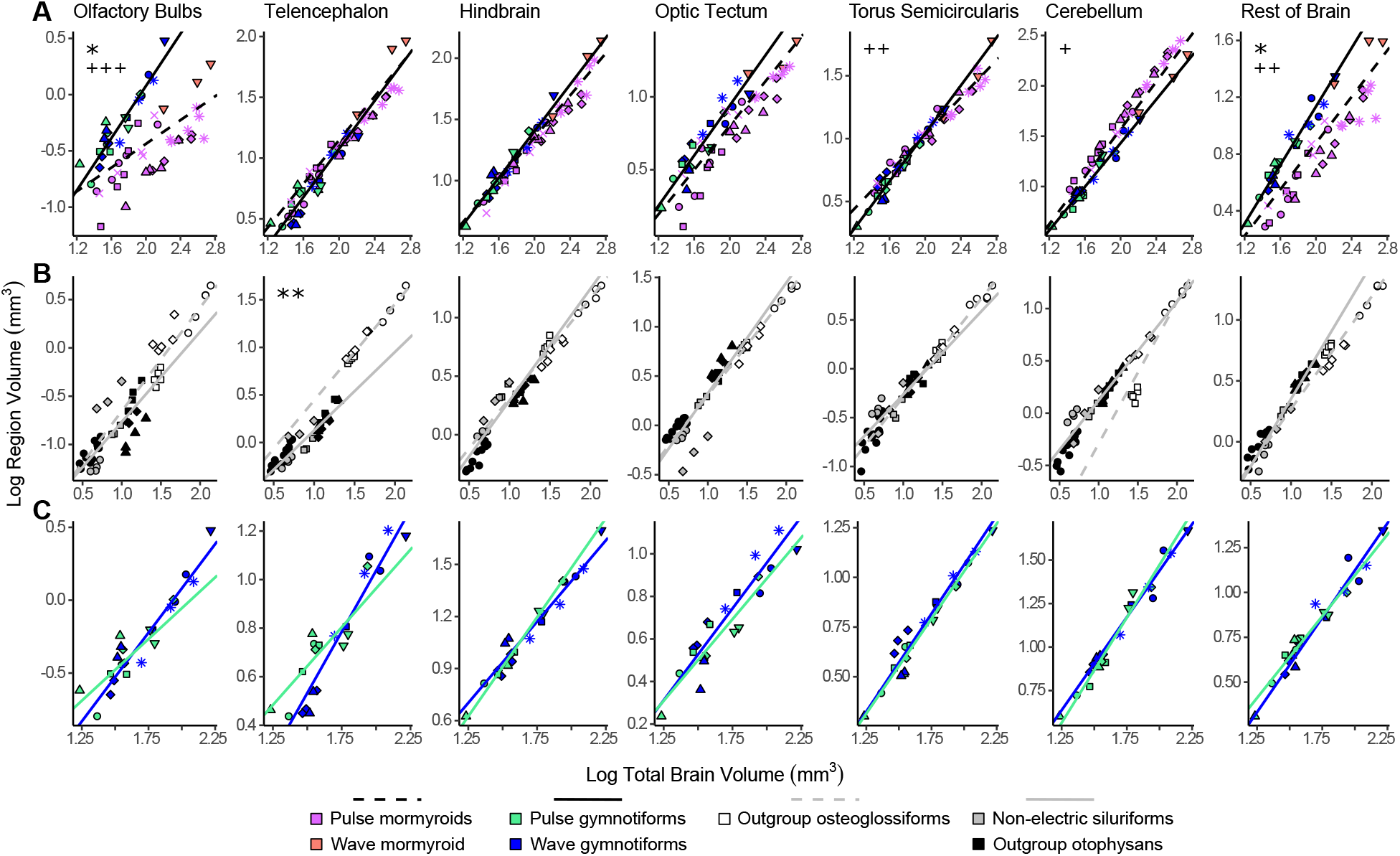
Lineage specific mosaic shifts within electrosensory phenotypes in olfactory bulbs, telencephalon, and rest of brain. Plots of log brain region volumes by log total brain volume for (A) mormyroids (pink and orange points, dashed black lines, N = 7) vs gymnotiforms (blue and green points, solid black line, N = 11); (B) non-electric osteoglossiforms (white points, dashed grey line, N = 3) vs non-electric otophysans (grey and black points, solid grey line, N = 7); and (C) wave (blue, N = 6) vs pulse (green, N = 5) gymnotiforms. Each point corresponds to an individual and shapes represent the same species as Figure 3. PGLS lines were determined from species means and compared using ANCOVAs. Significant differences in intercept are marked with asterisks: * = p < 0.05, ** = p < 0.01. Significant differences in slope are marked with pluses: + = p < 0.05, ++ = p < 0.01, +++ = p < 0.001. **Figure supplement 1.** ANCOVA results for lineage comparisons.

Some gymnotiforms produce continuous electrical discharges with variable frequency (wave-type), while others produce discrete electric discharges separated by variable periods of stasis (pulse-type). To assess whether there are mosaic shifts associated with evolution of electrical discharge type, we performed an ANCOVA between wave-type and pulse-type gymnotiforms (Figure 5C, Figure 5—figure supplement 1). We found no differences in either y-intercept or slope between wave and pulse gymnotiforms, suggesting that the transition between discharge types did not impact brain structure at this scale. It was not possible to directly test this in mormyroids since there is only one wave-type mormyroid species (*Gymnarchus niloticus*) that is sister to the family of all pulse-type mormyroids. However, the electrosensory system of the wave-type mormyroid is similar to that of gymnotiforms (Caputi et al., 2005), and the wave mormyroid trends more similar to gymnotiforms in both brain region residuals (Figure 5A) and the pPCA (Figure 3). After excluding the wave mormyroid, we found the same regional differences associated with all mormyroids, along with a significant increase in the cerebellum y-intercept in pulse mormyroids relative to gymnotiforms (p < 0.05) suggesting that the extraordinarily enlarged cerebellum of some mormyroid species is the result of both a steeper allometric relationship and a mosaic shift in the ancestor of pulse mormyroids.

## Discussion

We used osteoglossiform and otophysan fishes to test whether mosaic shifts in brain region volumes are associated with the convergent evolution of behaviorally novel active electrosensory systems. Although the mosaic shifts previously found in mormyroids were hypothesized to be related to the evolution of electrogenesis, it remained unknown if these patterns would be found in other electrogenic lineages. The brain scaling patterns of electrogenic and non-electric osteoglossiforms and otophysans are strikingly similar despite the considerable phylogenetic distance between them. Gymnotiforms have the most similar electrosensory system to mormyroids in terms of electrosensory structures, neural processing, and behavioral usage (Hopkins, 1995). In both of these lineages, we found mosaic increases in cerebellum, hindbrain, and torus semicircularis, which suggests that these evolutionary shifts in brain structure largely reflect their coevolution with electrogenesis and tuberous electroreceptor phenotypes. First-order electrosensory processing takes place in the electrosensory lateral line lobe (ELL) of the hindbrain, which projects to the torus for further processing of electrocommunication and electrolocation signals (Baker et al., 2013; Bell and Maler, 2005; Metzen and Chacron, 2021). The hindbrain is also involved in generating electromotor output (Caputi et al., 2005; Hagedorn et al., 1990; Kéver et al., 2020). Electrosensory information projects from the torus both directly and indirectly to areas of the cerebellum with large, reciprocal connections between cerebellum, torus, and ELL of both mormyroids and gymnotiforms. The overwhelming evidence of feedback circuits between initial electrosensory processing regions and cerebellum in both mormyroids and gymnotiforms suggests that the cerebellum may also be involved in processing electrosensory information in addition to motor control (Bell and Maler, 2005; Paulin, 1993). Cerebellum is known to be involved in predicting sensory consequences to motor movements and subsequent error detection, nonmotor functions, and learning (Hull, 2020; Popa and Ebner, 2019; Strick et al., 2009). Further, multiple areas of the mormyroid cerebellum show responses to electrosensory stimuli (Russell and Bell, 1978).

Electrogenic *Synodontis*, which only have ampullary electroreceptors, have significant mosaic increases in cerebellum and torus semicircularis relative to non-electric fishes. *Synodontis* electrical discharges are likely detectable by their ampullary electroreceptors (Hagedorn et al., 1990; Zupanc and Bullock, 2005) and involved in electrocommunication (Albert and Crampton, 2006; Boyle et al., 2014), which could relate to enlargement of these regions relative to electroreceptive but non-electric species. We found that *Synodontis* were intermediate between electrogenic fishes with tuberous receptors and non-electric fishes in torus and hindbrain, although the difference in hindbrain was not significant. The addition of tuberous electroreceptors increases the range of detectable signals and total electrosensory input to the brain relative to only ampullary electroreceptors (Crampton, 2019), and more subregions of the torus and hindbrain are devoted to electrosensory processing in tuberous electroreceptor species (Bell and Maler, 2005). The enlarged torus in *Synodontis* may also relate to acoustic communication. Some *Synodontis* species produce swim bladder sounds (Boyle et al., 2014), and the torus is involved in auditory processing (Fay and Edds-Walton, 2008). Interestingly, all electrogenic fishes have a significant mosaic increase in cerebellum regardless of electroreceptor type. Relative brain region sizes of electroreceptive only species are largely consistent with non-electrosensory species, which suggests that the evolution of electrogenesis has strong impacts on structural brain composition. However, two chondrichthyan lineages have independently evolved electrogenesis, Torpediniformes and Rajidae (Bennett, 1971), with no evidence of mosaic shifts in cerebellar size of these taxa (Mull et al., 2020; Yopak et al., 2010, Figure 4—figure supplement 3). It is possible this is because chondrichthyan cerebellums are already massively enlarged compared to their closest relatives, agnathans, who arguably lack a proper cerebellum (Striedter and Northcutt, 2020). Alternatively, it is possible the enlarged cerebellum in *Synodontis* is unrelated to the evolution of electrogenesis. Like *Synodontis*, Rajidae produce sporadic discharges likely used for electrocommunication, while mormyroids and gymnotiforms produce near continuous discharges (Bennett, 1971; Crampton, 2019). Considering we find a further enlargement in the cerebellum of pulse mormyroids who produce electrical discharges at varying and complex timing intervals, it is likely that the specific usage and complexities of the electrical discharges can influence the degree of cerebellar enlargement. We do not find any differences in cerebellar volume of wave and pulse gymnotiforms, but pulse gymnotiforms still produce discharges at regular intervals like wave-type fishes and unlike pulse mormyroids (Caputi et al., 2005). Further research on the usage of sporadic electrical discharges and the related electrosensory pathways are needed to better elucidate the relationship between electrogenesis and cerebellar enlargement.

Electrosensory information also projects to the telencephalon (Bell and Maler, 2005), but we do not find a mosaic increase in any electrosensory taxa. This is likely because telencephalon is involved in higher order sensory integration across many different sensory modalities (Striedter and Northcutt, 2020) while sensory systems remain segregated in the lower-order processing of hindbrain and torus (Meek and Nieuwenhuys, 1998). Surprisingly, we find that non-electric osteoglossiforms have a mosaic increase in telencephalon relative to non-electric otophysans. The telencephalon of osteoglossiforms is highly differentiated, more than other teleosts (Meek and Nieuwenhuys, 1998), which suggests that all osteoglossiforms may have had enlarged telencephalons that subsequently decreased in mormyroids alongside the evolution of electrogenesis and mosaic increases in other brain regions. Further research is needed to determine why these fishes have enlarged telencephalons. Optic tectum is also involved in sensorimotor integration, particularly with respect to the electrosensory, lateral line, and visual systems, in addition to being the primary target of visual input to the brain (Meek and Nieuwenhuys, 1998). Yet we find that tuberous receptor fishes have a mosaic decrease in optic tectum relative to non-electric fishes while electrogenic + ampullary only fishes are intermediate. Additionally, tuberous receptor fishes have a mosaic decrease in olfactory bulbs, which together could indicate decreased reliance on visual and olfactory systems. Gymnotiforms are thought to have poor vision (Takiyama et al., 2015), and different mormyroid lineages specialize to varying degrees in visual versus electrocommunication systems (Stevens et al., 2013). We do find a mosaic decrease in rest of brain in electrogenic taxa with tuberous electroreceptors and a mosaic decrease in rest of brain in mormyroids relative to gymnotiforms that could reflect a trade-off in one or more of the subregions that comprise the rest of brain, but since we combined these subregions, we are unable to speculate about their evolution.

Here, we assume that increased brain region volume corresponds to increased neuron number which increases processing power and reflects behavioral changes that natural selection can act upon. However, increases in regional volume can result from increased neuron number, neuron size, glia number, or any combination thereof. Previous studies found that neuron number and size tend to scale with brain size, but the degree varies both with lineage and brain region with some regions having more neurons than expected given total brain size (Barton, 2012; Herculano-Houzel et al., 2014; Marhounová et al., 2019). These findings suggest that volume measures could over- or underrepresent neuron number, and future studies should investigate neuron numbers in these regions to determine how region volumes are changing. Regardless of the specific differences in neuron numbers, we still find overwhelming evidence of differences in the volume of individual regions that, although the mechanism is currently unknown, are likely still reflecting biologically meaningful differences in neural processing across these fishes.

Different lineages can independently evolve the same phenotype via the same mechanism (parallel evolution) or different mechanisms (convergent evolution). Given the phylogenetic distance between osteoglossiforms and otophysans, it would be more remarkable to find that the different electrosensory systems and mosaic shifts in brain region volumes evolved in parallel rather than by convergence. Given that the mechanism of these regional increases remains unknown, we argue for a more conservative assumption of convergent evolution. This is supported by the fact that the cerebellar subregion that has expanded the most in mormyroids is the valvula cerebelli while in gymnotiforms, it is the corpus cerebelli (Meek and Nieuwenhuys, 1998). Additionally, the torus is laminar in gymnotiforms while there are distinct nuclei in the non-laminar torus of mormyroids (Bell and Maler, 2005). Regardless of whether the mechanism is convergent or parallel, we provide evidence of repeated, independent mosaic evolution of major brain regions in association with a convergent behavioral novelty. These findings demonstrate that evolutionary changes in gross-scale brain structure are surprisingly predictable alongside the evolution of active electrosensory systems. More broadly, these findings suggest that mosaic brain evolution may occur alongside the evolution of behavioral novelty and could reflect a degree of predictability in brain evolution with behavioral evolution.

## Materials and Methods

### Animal specimens

We measured structural brain variation for 63 individuals from 11 gymnotiform species, 7 siluriform species (4 from the electrogenic genus *Synodontis*), 2 characiform species, and 2 cypriniform species. Live cypriniforms, characiforms, siluriforms, and *E. virescens* were acquired through the aquarium trade and housed on a 12:12 light:dark cycle in 25-29°C water. Formalin-fixed gymnotiform and *S. petricola* specimens were provided by Dr. James Albert and Dr. Jason Gallant, respectively.

### Fixation

Live *Synodontis* were euthanized in 600 mg/mL tricaine methanesulfonate (MS-222) and immersion fixed in 4% buffered paraformaldehyde for two weeks. The remaining live fish were anesthetized in 300 mg/mL MS-222, euthanized by transcardial perfusion with 4% paraformaldehyde, and decapitated following methods in (Sukhum et al., 2018). These methods are consistent with euthanasia guidelines by the American Veterinary Medical Association and have been approved by the Animal Care and Use Committee at Washington University in St. Louis.

### *Synodontis* electrical recordings

Prior to fixation, we recorded from live *Synodontis* spp. following previous methods (Boyle et al., 2014) to determine whether they were electrogenic. Briefly, one or two individuals were placed into a tank containing a PVC tube for shelter and a differential recording electrode. We recorded continuously in two minute intervals for a total of 60 minutes. Signals were 500x amplified, bandpass filtered (1 Hz – 50 kHz, BMA-200, CWE Inc., Ardmore, PA), and digitized at 48.8 kHz (16-bit PCM converter, RX8, Tucker Davis Technologies, Alachua, FL) using custom MATLAB scripts. We recorded electrical discharges from all three tested species (Figure 6). We were unable to try recording from *S. petricola* to confirm electrogenesis. Lack of recording does not mean that they are incapable of producing electrical discharges, and only 1 out of 13 tested *Synodontis* species did not produce electrical discharges under experimental conditions (Baron et al., 1994, 2002; Boyle et al., 2014; Hagedorn et al., 1990). Further, electrogenic *Synodontis* spp. are not monophyletic and are broadly distributed throughout the species radiation (Day et al., 2013; Pinton et al., 2013), so additional research is needed to identify the number of origins and losses of electrogenesis among *Synodontis* fishes. Given the apparent lability of electrogenesis in *Synodontis* catfishes, these fishes would be good place to study the intermediate relationships between evolutionary changes in brain structure and evolution of electrogenesis.

**Figure 6.**
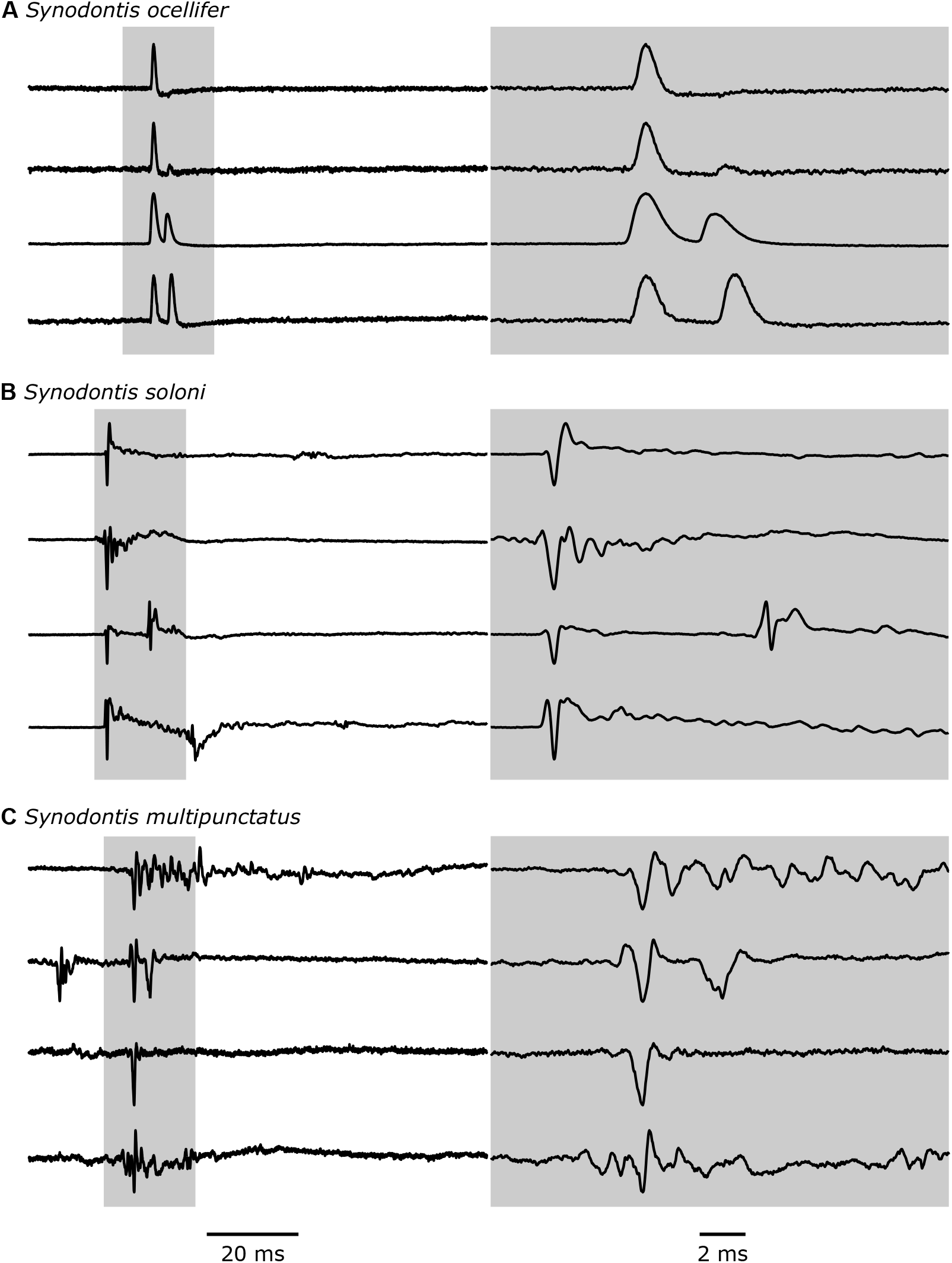
Electric discharges recorded from *Synodontis* spp. Four example traces that highlight the diversity of recorded electric discharges for *Synodontis ocellifer* (A), *Synodontis soloni* (B), and *Synodontis multipunctatus* (C). The left column shows traces with a scale bar of 20 ms. The shaded grey area is shown in the right column at a smaller time scale (scale bar = 2 ms). Discharges are amplitude-normalized and oriented in the same direction, but head-positive polarity is unknown.

### Micro-computed tomography scans

Heads were contrast stained in 5% phosphomolybdic acid (PMA) for one week for small specimens (mass < 14 g) or 8% PMA for two weeks for large specimens (mass ≥ 14 g) and then transferred to 0.1 M phosphate buffer. μCT scans were done at the Musculoskeletal Research Center at the Barnes-Jewish Institute of Health using a SCANCO μCT40 (Medical model 10 version SCANO_V1.2a, Brüttisellen, Switzerland) following scan conditions in (Sukhum et al., 2018). Slice thickness ranged from 6-18 μm, and scan tube diameter ranged from 12-36 mm depending on specimen size.

### Brain region delimitation

We used neuroanatomical landmarks to consistently delineate brain regions (Figure 7). The horizontal plane (white) divides the brain into dorsal and ventral areas. It extends from the most ventral point between the telencephalon and optic tectum (landmark a) to the most dorsal bulge of the spinal cord (landmark b).

**Figure 7.**
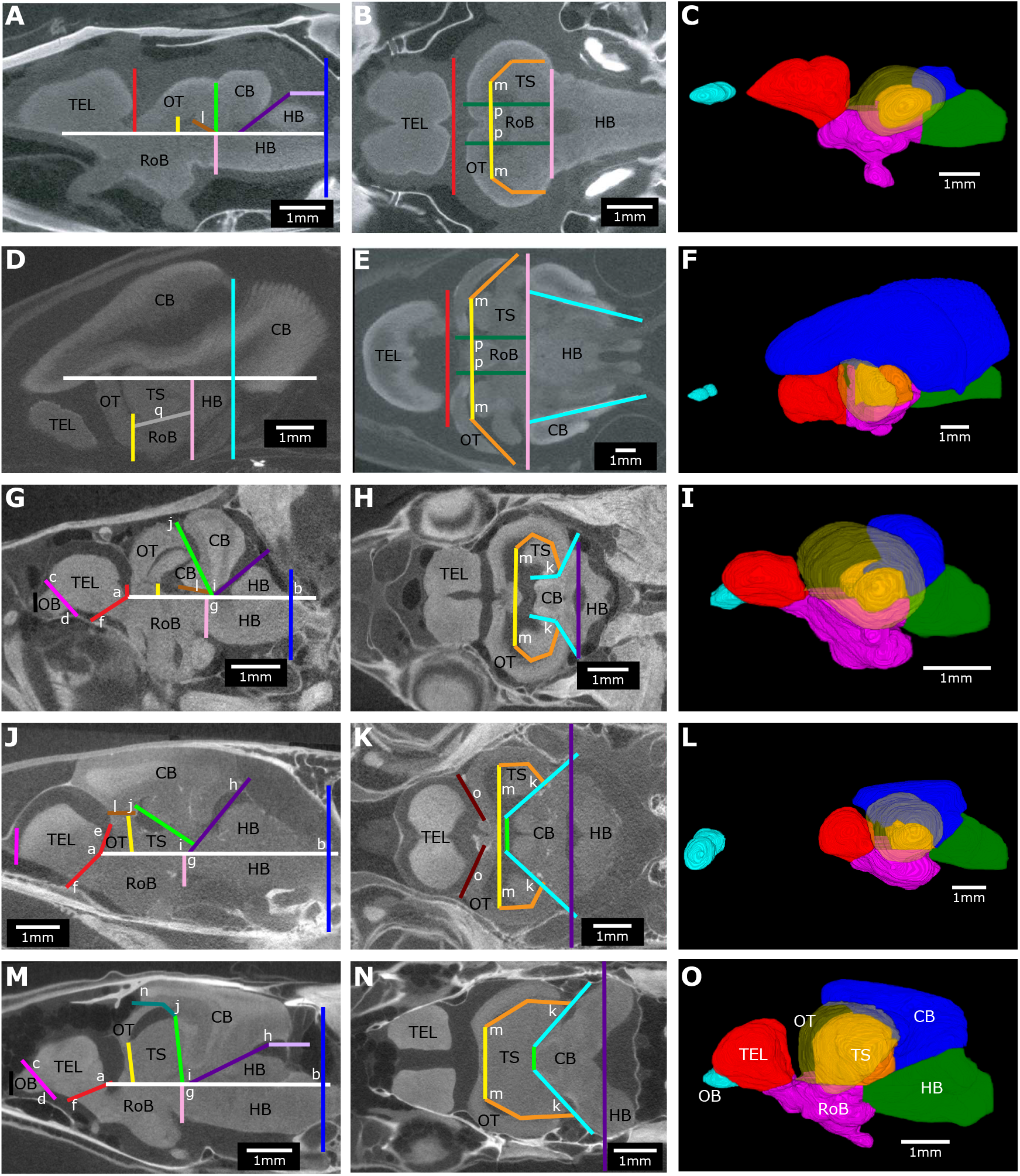
Brain landmarks and planes used to consistently delineate brain regions across species. Example brain slices and 3D reconstructions from *Pantodon buchholzi* (A-C), *Gnathonemus petersii* (D-F), *Phenocogrammus interruptus* (G-I), *Synodontis petricola* (J-L), and *Eigenmannia virescens* (M-O) that show the landmarks (letters) and planes (lines). Osteoglossiform brain slices (A, B, E) were modified from (Sukhum et al., 2018). Brain slices are oriented facing left in a sagittal plane (A, D, G, J, M) and horizontal plane (B, E, H, K, N). Images were made from averaging across ten slices (A, B, E) or five slices (D, G, H, J, K, M, N). 3D reconstructions have a semi-transparent optic tectum to show the torus semicircularis. Brain regions are color coded: OB = olfactory bulbs (cyan), TEL = telencephalon (red), HB = hindbrain (green), OT = optic tectum (yellow), TS = torus semicircularis (orange), CB = cerebellum (blue), RoB = rest of brain (magenta). Scale bar = 1 mm.

The olfactory bulb (OB) is a small, ellipsoid bulb at the anterior of the brain connected to the olfactory nerve. In gymnotiforms, characiforms, some cypriniforms, and some siluriforms, the olfactory bulb is the smaller bulb adjacent to the telencephalon. On the posterior side, the olfactory bulb is separated by the olfactory plane (magenta), which starts at the most posterior point of the anterior side of the telencephalon (landmark c) and extends to the most dorsal point underneath the telencephalon (landmark d). On the anterior side, the olfactory bulb is separated from the olfactory tract by a straight plane (black) at the base of the bulge that is the olfactory bulb. In the remaining species, the olfactory bulb is in the anterior region of the skull cavity and is clearly separated from the rest of the brain by the olfactory nerve.

The telencephalon (TEL) is the larger ellipsoid bulb at the anterior of the brain. There is a clear fissure separating the telencephalon from the more posterior regions of the brain. On the anterior side, the telencephalon is separated by the olfactory plane (magenta). The posterior-TEL plane (red) is a connection of three points: a. the most ventral point between the telencephalon and optic tectum, where it meets the horizontal plane, e. the most posterior bulge of the telencephalon, and f. the lower concave curve of the telencephalon, which is just anterior to the optic nerve. In some cases, a and e are the same point.

The hindbrain (HB) is the most posterior region both above and below the horizontal plane. On the anterior side above the horizontal plane, the plane separating the hindbrain (CB-HB plane, dark purple) extends from the most ventral point between the hindbrain and cerebellum (landmark g) to the concave curve of the hindbrain (landmark h). This most ventral point is a clear cistern separating the cerebellum and hindbrain when viewed in a frontal slice. In species where the cerebellum extends dorsally over the hindbrain, the horizontal-CB plane (light purple) extends from the end of the CB-HB plane (dark purple) parallel to the horizontal plane along the ventral side of the cerebellum. On the anterior side below the horizontal plane, the hindbrain is separated by the anterior-HB plane (light pink) marked by the concave curve of the cerebellum (landmark i) and extending in a straight line perpendicular to the horizontal plane (white). On the posterior side, the hindbrain is separated from the spinal cord by the posterior-HB plane (dark blue): a straight line perpendicular to the horizontal plane that is marked by the most posterior point of the cerebellum or dorsal bulge of the spinal cord (landmark b), whichever is most posterior.

The cerebellum (CB) is the most dorsal region of the brain. It extends from the optic tectum to the hindbrain, but sometimes covers the telencephalon in mormyroids. On the anterior side, the cerebellum is clearly separated from the optic tectum and torus semicircularis. Following this separation, the anterior-CB plane (light green) extends from the top of the optic tectum (landmark j) to the horizontal plane. The lateral-CB plane (cyan) separates the remainder of the torus semicircularis from the cerebellum and connects from the end of the anterior-CB plane and follows the posterior curve of the torus semicircularis to connect to the most posterior concave curve of the torus semicircularis (landmark k). On the posterior side, the cerebellum is separated from the hindbrain by the CB-HB plane (purple). On the ventral side, the cerebellum is separated from the hindbrain and rest of brain by the horizontal plane (white). In *Synodontis*, the optic tectum and torus semicircularis are more lateral and the anterior end of the cerebellum extends further ventral. To separate this part of the cerebellum from the optic tectum, there is an additional ventral-CB plane (brown) extending between the posterior-TEL plane (red) and OT-TS plane (yellow) along the most ventral point at the anterior of the cerebellum (landmark l). In non-electric species, the cerebellum extends anteriorly between the optic tectum and dorsal to the torus semicircularis and rest of brain.

To define these anterior boundaries of the cerebellum, the lateral-CB plane (cyan) consists of a second plane that extends further anterior along the lateral sides of the cerebellum. The ventral-CB plane (brown) extends between the most anterior point of the cerebellum to the anterior-CB plane (light green) along the most ventral point at the anterior of the cerebellum (landmark l).

The optic tectum (OT) is the most lateral bulge of the brain and forms a cup like structure around the torus semicircularis and rest of the midbrain. On the anterior side, the optic tectum is separated from the telencephalon by the posterior-TEL plane (red). On the lateral and posterior sides, the optic tectum is separated from the torus semicircularis by the OT-TS plane (yellow) and the lateral-TS planes (orange). The OT-TS plane (yellow) follows the curve of the torus semicircularis and extends medial-laterally connecting the furthest anterior curves of the torus semicircularis (landmark m). The lateral-TS planes (orange) extend from the end of the OT-TS plane (yellow) to the furthest lateral curve of the torus semicircularis. In gymnotiforms, this requires two planes, but in siluriforms, characiforms, and cypriniforms, this requires three or four planes due to the optic tectum wrapping more tightly around the torus semicircularis. Dorsally, the optic tectum is separated from the cerebellum by the OT-CB plane (teal), which extends from the end of the anterior-CB plane (light green) following along the curve of the optic tectum to the most anterior, concave curve of the cerebellum (landmark n). This requires two planes in some species due to a more anteriorly extended cerebellum. In *Synodontis*, the optic tectum is more distal to the midline of the brain than in gymnotiforms and instead the OT-CB plane (teal) extends from the most medial and ventral point separating the optic tectum from the cerebellum along the curve of the optic tectum to the anterior-CB plane (light green). In *Synodontis*, there is an additional plane to separate the anterior of the optic tectum from the cerebellum; this anterior-OT plane (dark red) extends from the most anterior curve of optic tectum (landmark o) moving medially along the curve of the optic tectum to the OT-CB plane (teal).

The torus semicircularis (TS) is the two symmetrical, ellipsoid bulbs within the cup of the optic tectum. The torus semicircularis is clearly separated from the more anterior and lateral optic tectum by the OT-TS plane (yellow) and the lateral-TS planes (orange). On the posterior side, the torus semicircularis is clearly separated from the cerebellum by the anterior-CB plane (light green), lateral-CB plane (cyan), and ventral-CB plane (brown). On the ventral side, the torus semicircularis is separated from the rest of brain by the horizontal plane (white). For the osteoglossiform brains, we used the landmarks and planes in (Sukhum et al., 2018) with additional planes to separate the torus semicircularis from the rest of brain. The boundaries of the torus semicircularis in outgroup osteoglossiforms are equivalent to those used for outgroup otophysans with the addition of a medial boundary (optic tectum medial planes, dark green) to separate the torus semicircularis from the rest of brain. The optic tectum medial planes (dark green) extend along the furthest lateral curve of the thalamus (landmark p) as in (Sukhum et al., 2018) but were modified to extend further posterior to intersect the anterior-HB plane (light pink). In mormyroids, the enlarged cerebellum pushes the torus semicircularis further ventral, below the horizontal plane (white). The torus semicircularis is separated from the optic tectum by the OT-TS plane (called optic tectum plane in (Sukhum et al., 2018), yellow) and the lateral-TS planes (lateral optic tectum planes, orange). The torus semicircularis is separated dorsally from the cerebellum by the horizontal plane (white), posteriorly from the hindbrain by the anterior-HB plane (light pink), and medially from the rest of brain by the optic tectum medial planes (dark green) which were modified to extend further posterior to the anterior-HB plane (light pink). On the ventral side, the torus semicircularis is separated from the rest of brain by the ventral-TS plane (grey), which extends from the most ventral point between the torus semicircularis and optic tectum along the most ventral curve of the torus semicircularis (landmark q) to the anterior-HB plane (light pink).

The rest of brain (RoB) combines the remainder of the undifferentiated brain into one region and is between the horizontal plane (white), posterior-TEL plane (red), and anterior-HB plane (light pink).

We followed the natural breaks in continuous brain tissue wherever possible, which on occasion permitted continuous brain tissue to cross the boundaries set by the planes. We only allowed this when the natural breaks in the brain tissue were obvious and unambiguous, and we never allowed for crossing of the posterior-HB plane (dark blue).

In (Sukhum et al., 2018), the authors did not separate torus semicircularis from rest of brain because the area that they had defined as torus semicircularis also included non-toral regions of the midbrain. Here, we have decided to separate torus semicircularis from rest of brain despite this because the non-toral regions of the midbrain that are included within the boundaries of our definition of torus semicircularis are comparable across all of our species with the largest non-toral regions included in the torus semicircularis of our non-electrosensory species. This means that although the torus semicircularis volume is overestimated some in all species, the overestimation is larger in our non-electrosensory species than in our electrosensory species, and thus our findings are potentially more conservative than the real differences in regional volumes. Further, the absolute region volumes are not the focus of the study, rather we are concerned with the relative patterns of region volumes across taxa. A consistent overestimation of torus semicircularis volumes does not change these patterns.

### Quantifying region volume

Brain region volumes were measured using the ImageJ Volumest plugin (Merzin, 2008; Schindelin et al., 2012; Schneider et al., 2012). We manually traced each region every 2-10 slices: regions < 2 mm^3^ were measured every 2 slices, regions 2-4 mm^3^ were measured every 5 slices or less, and regions > 4 mm^3^ were measured every 10 slices or less. Volumest then calculates volume using stereological methods with slice thickness ranging from 6-18 μm, depending on specimen size, and a 0.1 mm grid width. We randomly selected 4 scans to be remeasured twice, blind to previous results and species identity. Coefficients of variation for these remeasures were all less than 4% (Supplementary Table 1).

### Phylogenetic analysis

We built a Bayesian phylogenetic tree from 6 aligned and concatenated genes (*16s*, *col*, *cytb*, *rh1*, *rag1*, and *rag2*) of 189 species spanning Anguilliformes to Ostariophysi using Beast v1.10.4 (Suchard et al., 2018). We used a birth-death process tree prior and unlinked, relaxed lognormal clock models. We used unlinked substitution models of HKY+I+G for *rh1* and GTR+I+G for all other genes as determined by jModelTest (Darriba et al., 2012; Guindon and Gascuel, 2003). To reduce the computational burden and improve taxonomic resolution, we constrained the monophyly of each order, gymnotiform families, elopomorpha, ostariophysi, and gymnotiforms + siluriforms as sister to each other following previous studies that used substantially more sequence data and tested alternative hypotheses of teleost topology but did not include all of the species used in this study (Hughes et al., 2018; Rabosky et al., 2013). We time calibrated the phylogeny using the fossil dates and justifications in (Rabosky et al., 2013). We performed two independent Bayesian analyses starting from random trees each with a chain length of 150,000,000 sampled every 10,000 generations. We used Tracer v1.7.1 to confirm convergence of parameter values across both analyses and effective sample size values > 200 (Rambaut et al., 2018). We combined the output of both analyses after discarding the first 15,000,000 states for each run and estimated the maximum clade credibility tree. To include data from unsequenced species, we used sequence data from the species in the same monophyletic genus with the shortest distance to the genus node. We pruned the tree to only include species with brain measurement data. Given the uncertainty of phylogenetic relationships both among otophysans and within gymnotiforms (Crampton, 2019; Hughes et al., 2018; Rabosky et al., 2013) and our use of relatively few previously sequenced genes, we make no claims that these are the actual phylogenetic relationships between these species. Instead our purpose in this study was to correct for relatedness as best we could.

We performed a phylogenetic principal components analysis (pPCA) on species means and applied those rotations to the data from all individuals. We compared phylogenetic generalized least squares (PGLS) models considering the null hypothesis of body mass and total brain volume predicting PC1-4 values to PGLS models considering that and either and both of the electrosensory phenotypes. Since *G. niloticus* individuals were received by (Sukhum et al., 2018) as fixed, decapitated specimens, their body mass was unknown, thus we removed them from all PC model fits. PC1 was correlated with total brain volume, as expected in allometric relationships (Klingenberg, 1996), so we removed total brain volume as a covariate in all PC1 models. We allowed the strength of phylogenetic signal to vary in each model using Pagel’s lambda (λ) where 0 indicates no phylogenetic signal and 1 indicates phylogenetic signal consistent with Brownian motion (Pagel, 1999). All models were fit following maximum likelihood and compared using small-sample corrected Akaike information criterion (AICc) with a ΔAICc cutoff of 2 (Burnham and Anderson, 2002). All PGLS fits were determined using species means.

To test for mosaic shifts, we fit PGLS regressions of each brain region volume and total brain volume for each group. We allowed λ to vary for each brain region and tested for significant differences between groups using an analysis of covariance (ANCOVA). For the three electrosensory phenotypes, we performed pairwise post hoc tests with a Bonferroni correction for each region with significant differences. To determine relative brain sizes, we performed a PGLS regression of total brain volume against body mass for all species excluding *G. niloticus*. All phylogenetic analyses were done using R v3.6.2 and the packages phytools, ape, nlme, MuMIn, and emmeans (R Core Team, 2019; Lenth, 2020; Bartoń, 2020; Pinheiro et al., 2019; Revell, 2012; Paradis et al., 2004).

## Acknowledgments

We thank James Albert for providing additional gymnotiform specimens, Jason Gallant for providing *Synodontis* spp., and Emilia Martins, Piyumika Suriyampola, and Anuradha Bhat for providing wild-caught *D. rerio*. This work was funded by NSF IOS-1755071 to B.A.C.

## Author Contributions

E.L.S. and B.A.C. designed research; E.L.S. performed research; E.L.S. analyzed data; E.L.S. and B.A.C. wrote the paper.

## Figure Supplements

**Figure 1—figure supplement 1 (separate file).** Example 3D reconstructions of brains from Figure 1*B*. Brains are oriented from a lateral view with anterior to the left and dorsal at the top. Brain regions are color coded: OB = olfactory bulbs (cyan), TEL = telencephalon (red), HB = hindbrain (green), OT = optic tectum (yellow), TS = torus semicircularis (orange), CB = cerebellum (blue), RoB = rest of brain (magenta). Scale bar = 1 mm.

**Figure 3—figure supplement 1.**
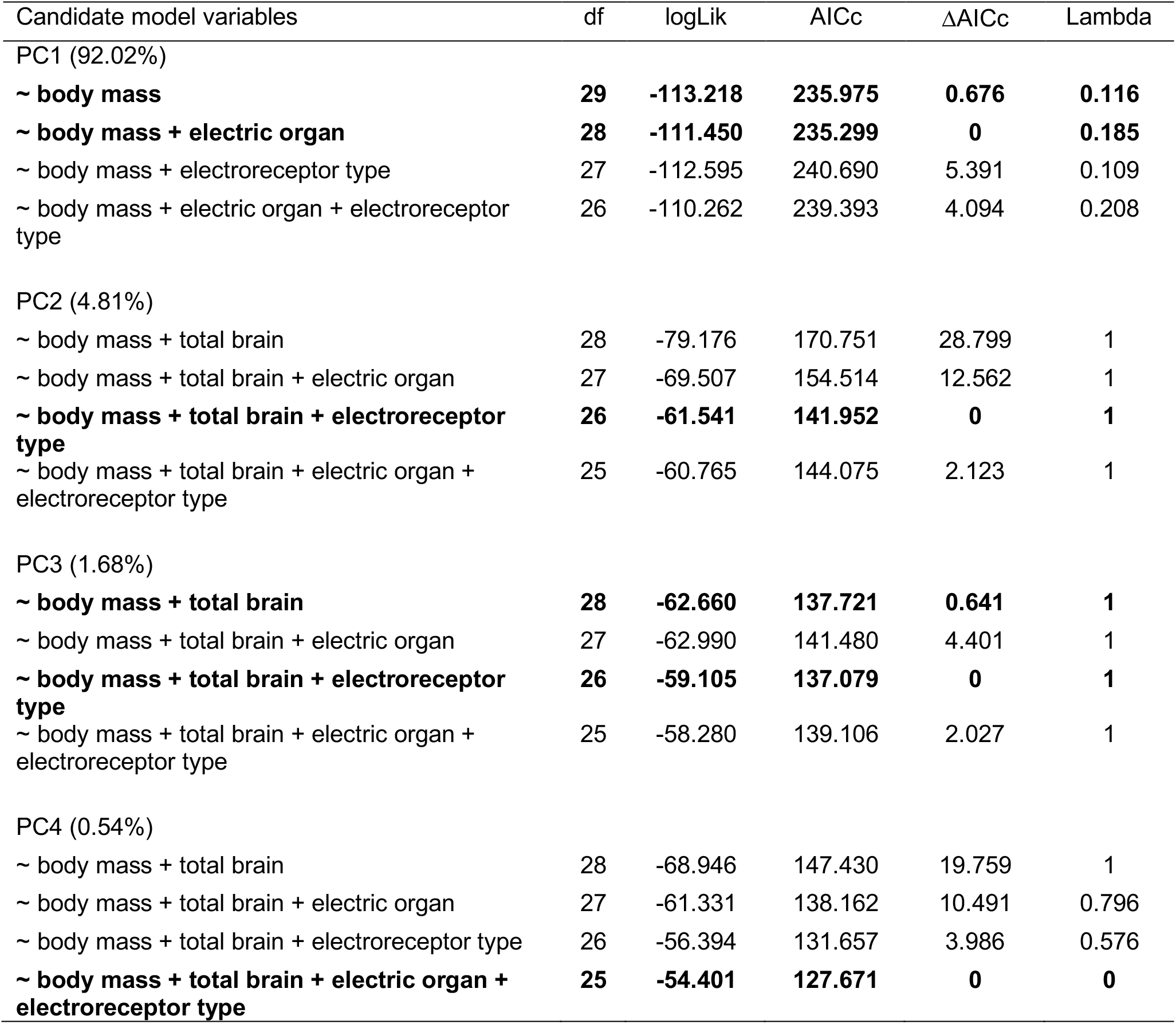
pPCA PGLS model selection results (N = 31). PC1 was correlated with total brain volume, thus total brain volume was removed from all PC1 models.

**Figure 4—figure supplement 1.**
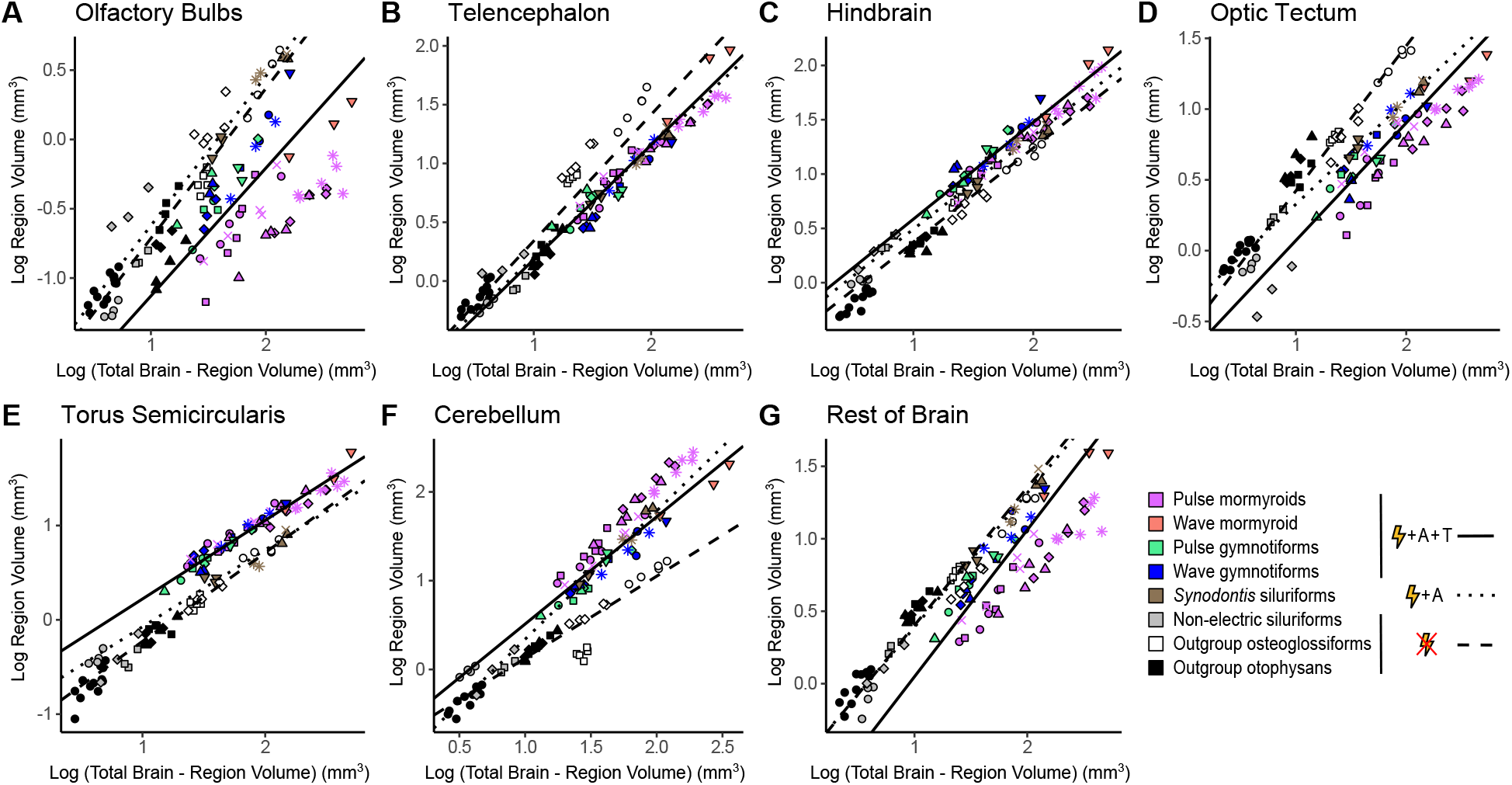
Apparent mosaic shifts in olfactory bulbs, hindbrain, optic tectum, torus semicircularis, cerebellum, and rest of brain between electrogenic species with ampullary & tuberous electroreceptors and non-electric species. Plots of log region volume against log total brain – region volume for olfactory bulbs (A), telencephalon (B), hindbrain (C), optic tectum (D), torus semicircularis (E), cerebellum (F), and rest of brain (G). Each point corresponds to an individual and shapes represent the same species as Figure 3. PGLS lines were determined from species means and correspond to electrosensory phenotypes that cluster distinctly in Figure 3: electrogenic + tuberous & ampullary electroreceptors (pink, orange, green, blue points; solid line; N = 18), electrogenic + only ampullary electroreceptors (brown points; dotted line; N = 4), and non-electric (grey, white, black points; dashed line; N = 10).

**Figure 4—figure supplement 2.**
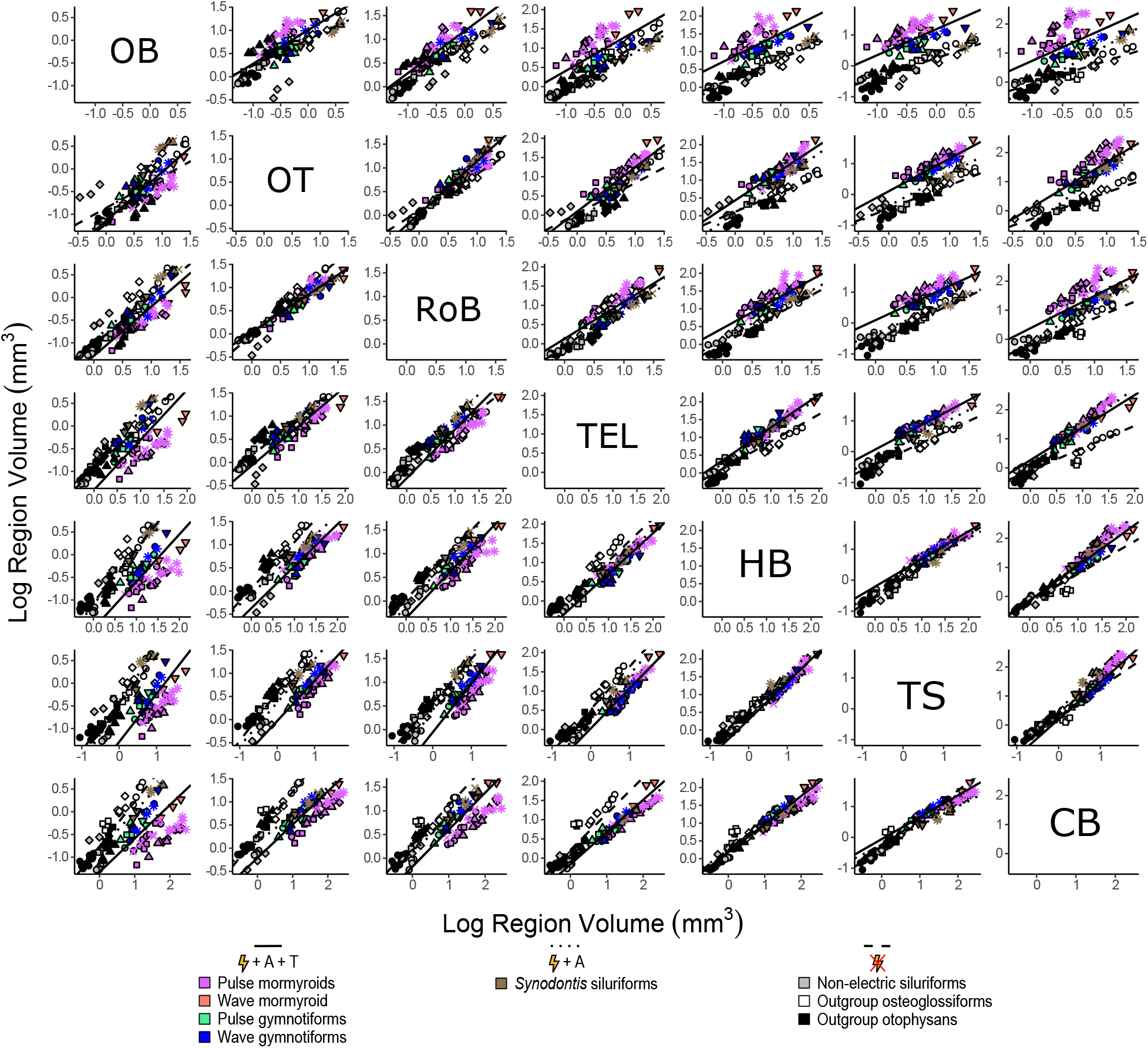
Apparent mosaic shifts in y-intercept between electrogenic species with ampullary & tuberous electroreceptors and non-electric species for many region to region comparisons. Matrix of scatterplots for each region by region comparison. Each point corresponds to an individual and shapes represent the same species as Figure 3. PGLS lines were determined from species means and correspond to electrosensory phenotypes that cluster distinctly in Figure 3: electrogenic + tuberous & ampullary electroreceptors (pink, orange, green, blue points; solid line; N = 18), electrogenic + only ampullary electroreceptors (brown points; dotted line; N = 4), and non-electric (grey, white, black points; dashed line; N = 10).

**Figure 4—figure supplement 3.**
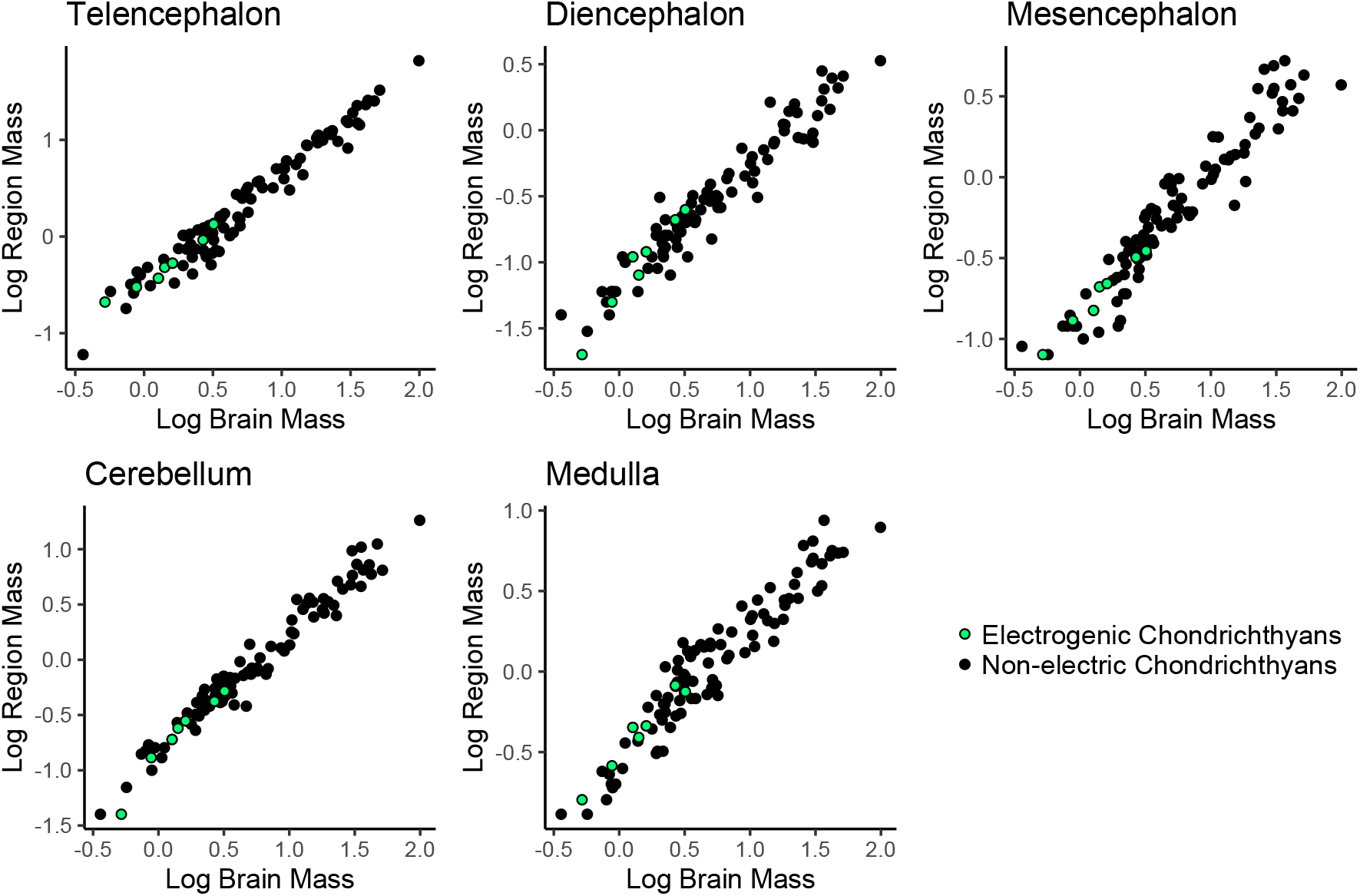
No evidence of mosaic shifts in cerebellum or medulla of electrogenic chondrichthyans. Plots of log region mass against total brain mass, data from (Mull et al., 2020). Each point is a species, electrogenic taxa are in green, and non-electric taxa are in black.

**Figure 4—figure supplement 4.**
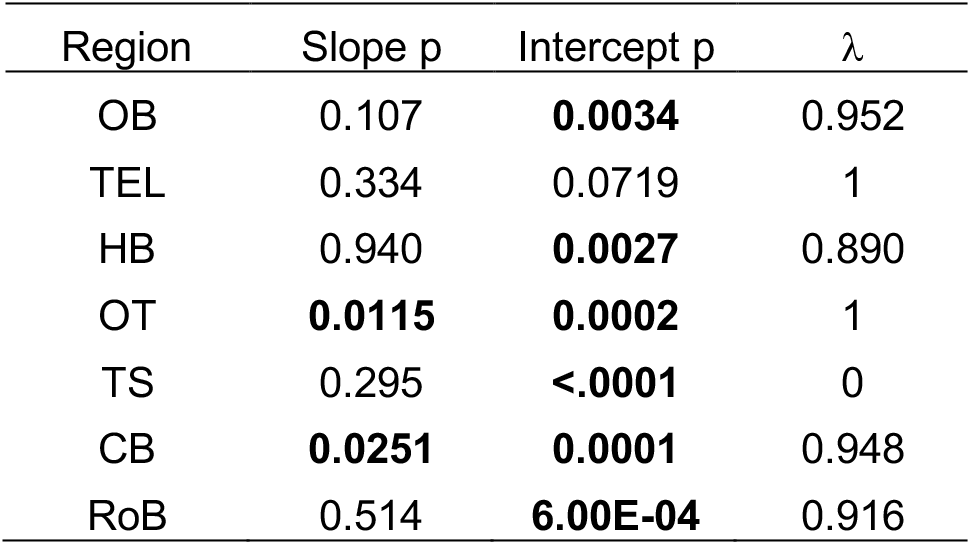
Results of an ANCOVA comparing PGLS relationships of region volume against total brain volume for each electrosensory phenotype: slope p, intercept p, and Pagel’s lambda (λ). OB = olfactory bulbs, TEL = telencephalon, HB = hindbrain, OT = optic tectum, TS = torus semicircularis, CB = cerebellum, RoB = rest of brain.

**Figure 4—figure supplement 5.**
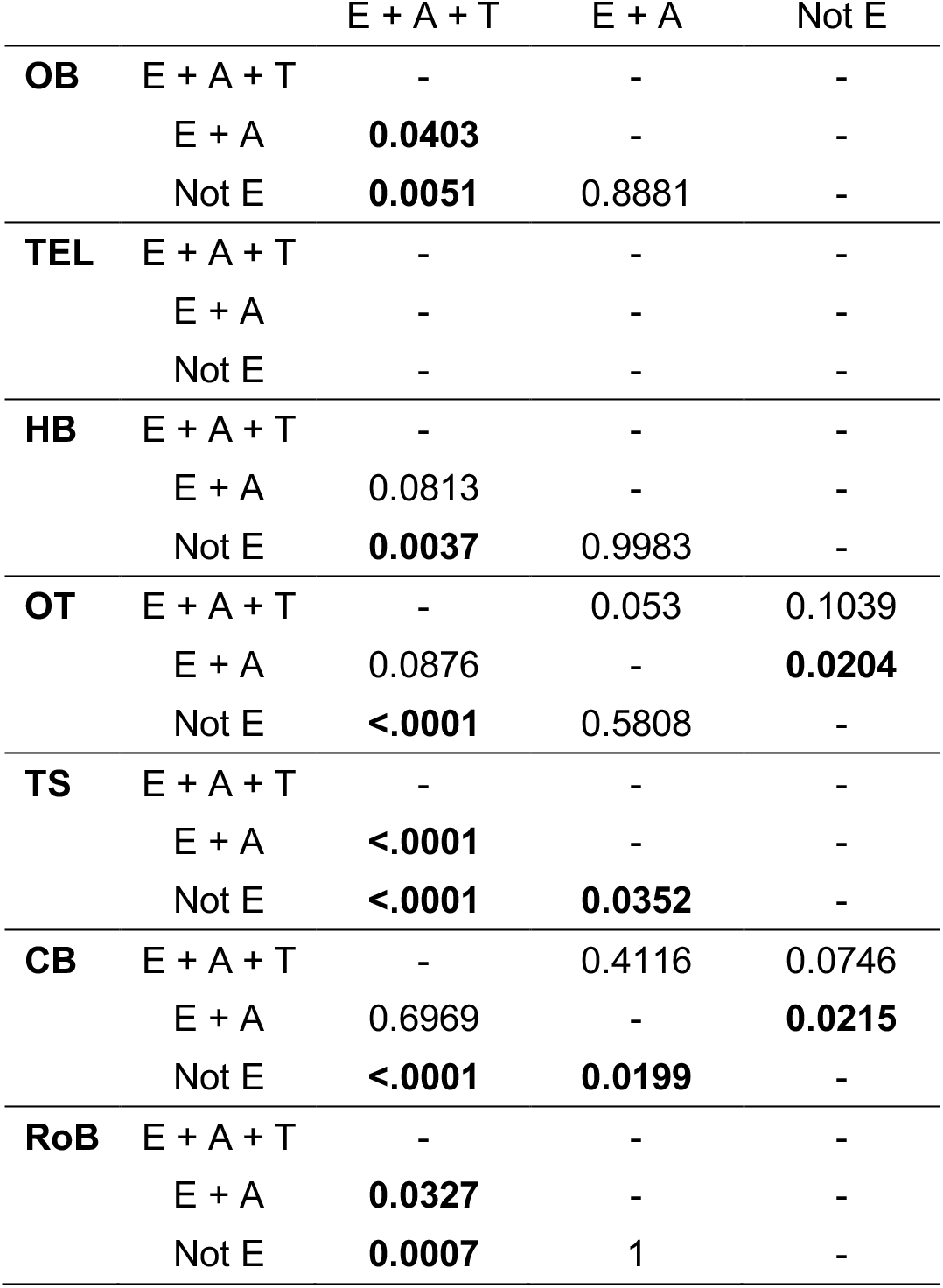
Results (p values) of pairwise posthoc tests with a Bonferroni correction for an ANCOVA comparing PGLS relationships of region volume against total brain volume for each electrosensory phenotype: electrogenic + tuberous & ampullary electroreceptors (E + A + T), electrogenic + only ampullary electroreceptors (E + A), and non-electric (Not E). Differences in intercept are below the diagonal and differences in slope are above the diagonal for each brain region. OB = olfactory bulbs, TEL = telencephalon, HB = hindbrain, OT = optic tectum, TS = torus semicircularis, CB = cerebellum, RoB = rest of brain.

**Figure 4—figure supplement 6.**
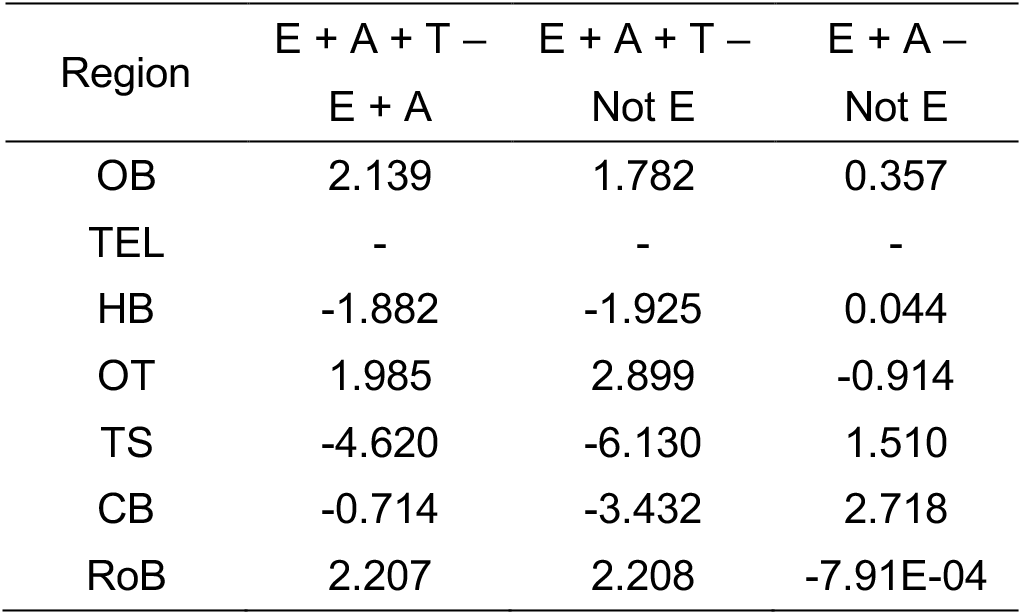
Estimated effect sizes (Cohen’s d) of each contrast for an ANCOVA comparing PGLS relationships of region volume against total brain volume for each electrosensory phenotype: electrogenic + tuberous & ampullary electroreceptors (E + A + T), electrogenic + only ampullary electroreceptors (E + A), and non-electric (Not E). OB = olfactory bulbs, TEL = telencephalon, HB = hindbrain, OT = optic tectum, TS = torus semicircularis, CB = cerebellum, RoB = rest of brain.

**Figure 5—figure supplement 1.**
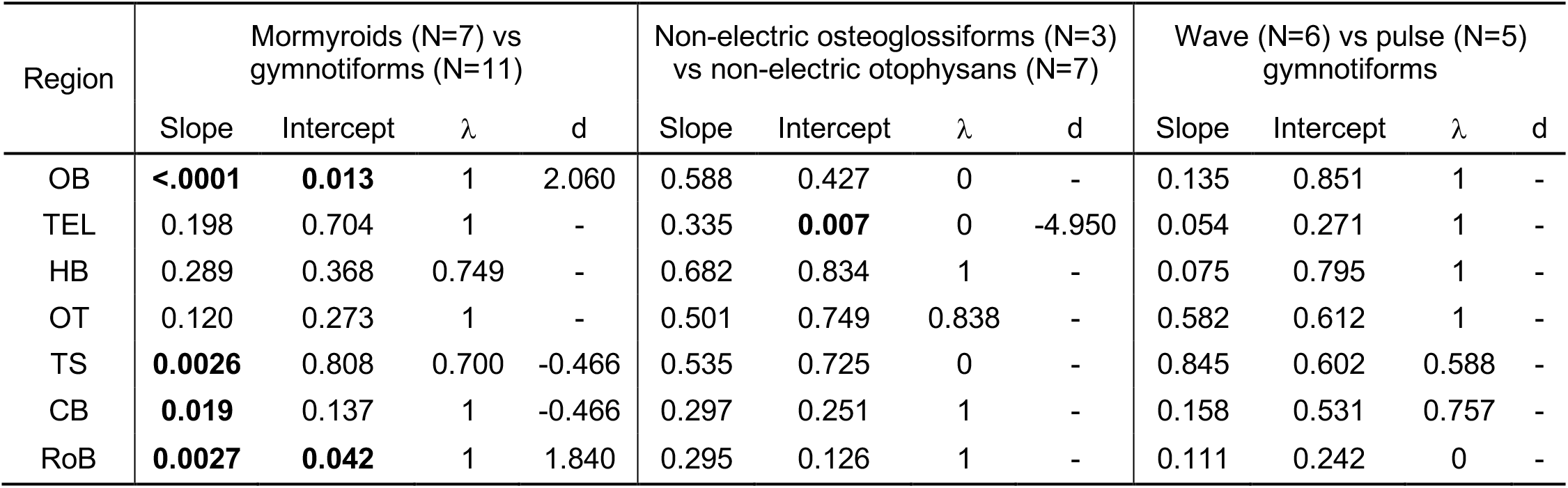
Lineage ANCOVA results comparing PGLS relationships of region volume against total brain volume for each lineage: slope p, intercept p, Pagel’s lambda (λ), Cohen’s d. OB = olfactory bulbs, TEL = telencephalon, HB = hindbrain, OT = optic tectum, TS = torus semicircularis, CB = cerebellum, RoB = rest of brain.

**Supplementary Table 1.** Coefficient of variation results of repeated measures (N = 3) for four different brains, related to Materials and Methods. OB = olfactory bulbs, TEL = telencephalon, HB = hindbrain, OT = optic tectum, TS = torus semicircularis, CB = cerebellum, RoB = rest of brain, TBV = total brain volume.

| Species | OB | TEL | HB | OT | TS | CB | RoB | TBV |
| --- | --- | --- | --- | --- | --- | --- | --- | --- |
| *S. elegans* | 0.95% | 0.95% | 1.35% | 1.13% | 1.32% | 1.31% | 2.74% | 1.24% |
| *S. macrurus* | 1.35% | 0.54% | 2.16% | 1.53% | 0.06% | 1.16% | 2.16% | 0.61% |
| *S. multipunctatus* | 3.97% | 2.72% | 3.86% | 0.51% | 3.53% | 0.28% | 3.21% | 0.28% |
| *P. interruptus* | 0.70% | 0.45% | 1.65% | 0.42% | 1.92% | 0.67% | 2.39% | 0.20% |

**Supplementary file 1 (separate file).** Brain region data from this study. OB = olfactory bulbs, TEL = telencephalon, HB = hindbrain, OT = optic tectum, TS = torus semicircularis, CB = cerebellum, RoB = rest of brain.

## Notes

### Competing Interest Statement

The authors have declared no competing interest.

